# Neural Basis of The Double Drift Illusion

**DOI:** 10.1101/2022.01.25.477714

**Authors:** Noah J. Steinberg, Zvi N. Roth, J. Anthony Movshon, Elisha P. Merriam

**Author notes:** These authors contributed equally to this work (Noah J. Steinberg, Zvi N. Roth). Correspondence should be addressed to N.J.S. and Z.N.R. Author contributions: EPM and JAM conceived and designed the experiment. NJS and EPM conducted the experiment. NJS, ZNR, and EPM collected and analyzed the data, and wrote and revised the manuscript.

## Abstract

In the “double-drift” illusion, local motion within a window moving in the periphery alters the window’s perceived path. The illusion is strong even when the eyes track a target whose motion matches the window so that the stimulus remains stable on the retina. This implies that the illusion involves the integration of retinal signals with non-retinal eye-movement signals. To identify where in the brain this integration occurs, we measured BOLD fMRI responses in visual cortex while subjects experienced the double drift illusion. We identified a number of cortical areas that responded more strongly during the illusion, but only in area hMT+ was it possible to decode the illusory trajectory. Our results provide evidence for a perceptual representation in human visual cortex that is not linked to retinal position.

## Introduction

The primate visual system is retinotopic: neurons throughout visual cortex encode the location of visual stimuli on the retina (Gardner et al., 2008). Yet visual perception is stable across frequent eye movements that displace the retinal image. This observation has led to the conjecture that the brain maintains a world-centered or ‘spatiotopic’ representation that is invariant to changes in eye position (Duhamel et al., 1997).

A number of human imaging studies have reported ‘spatiotopic’ responses in human visual cortical areas, including lateral occipital complex (LOC), the human analogue of MT and MST (hMT+), and area V6 (d’Avossa et al., 2007; McKyton and Zohary, 2007; Crespi et al., 2011). However, other studies have failed to find such evidence (Gardner et al., 2008; Golomb and Kanwisher, 2012; Merriam et al., 2013), and this discrepancy has not been fully resolved. One suggestion is that the reference frame for stimulus encoding depends on cognitive or task demands. For example, Crespi et al. (2011) reported that the reference frame of visual responses can shift from retinotopic to spatiotopic, depending on the attentional state of the observer. Behavioral studies have reported that spatiotopic representations become more prominent in tasks requiring sequences of eye movements, suggesting that world-centered coordinates are built-up over time (Poletti et al., 2013; Sun and Goldberg, 2016).

Here, we used a version of the double-drift illusion to investigate spatiotopic processing. The double drift illusion is a particularly striking phenomenon because of the magnitude of the illusion: a combination of local horizontal motion with a global vertical motion trajectory causes a strong perception of an illusory drift away from the veridical trajectory that can exceed 45 deg (Tse and Hsieh, 2006; Shapiro et al., 2010; Lisi and Cavanagh, 2015). A recent study revealed that the illusion persists even during smooth pursuit when the stimulus is stabilized on the retina (Cavanagh and Tse, 2019).

The pursuit version of the double drift illusion could provide insight into spatiotopic encoding in the brain. The fact that the illusion survives smooth pursuit when the stimulus is in fact stable on the retina suggests that the trajectory of the illusion is encoded in non-retinal coordinates. We hypothesized that several regions in occipital and parietal cortex are candidate regions involved in the illusory percept.

Many of these areas encode stable stimulus position during pursuit eye movements (i.e. ‘real position’ cells) (Nau et al., 2018). Moreover, several of these areas have been implicated in spatiotopic processing (d’Avossa et al., 2007). If our hypothesis is correct, we predict that activity in extrastriate cortex will reflect the illusory motion path instead of the veridical stimulus path.

## Results

We tested whether fMRI BOLD activity in human visual cortex reflects the perceived spatial position of a visual stimulus that remained at a constant retinal location. We measured BOLD activity during a version of the double drift illusion in which the perceived location of the stimulus differed from its actual location by several degrees (Fig 1).

**Figure 1:**
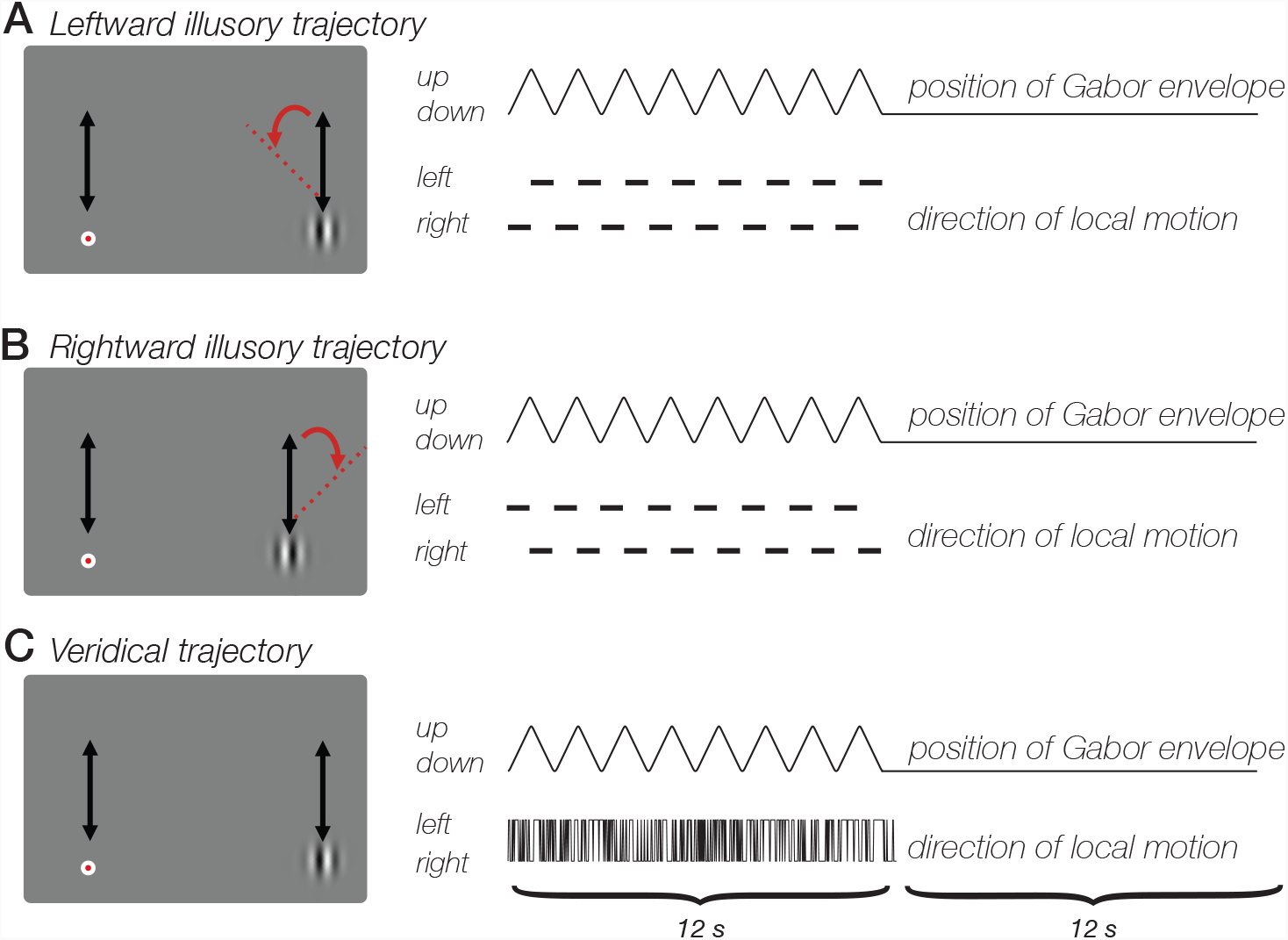
Double drift illusion during smooth pursuit. **Top**, Leftward drift illusion. Participants made smooth pursuit eye movements, tracking the red dot as it moved vertically in tandem with a Gabor stimulus. Conjunction of local motion (grating phase drift) and global motion (displacement of the Gaussian envelope) produces an illusion in which the Gabor appears to drift several degrees to the left of its actual trajectory, even when smooth pursuit eye movements stabilize the Gabor on the retina. **Middle**, Rightward drift illusion. Conjunction of local and global motion produces strong illusion of a rightward Gabor trajectory. **Bottom**. No-illusion control condition. Randomly-updated grating phase does not produce illusory stimulus trajectory. All three stimulus conditions contain the same net motion energy and involved the same pursuit eye movements, yet are associated with strongly different percepts.

To determine whether BOLD activity contained information about the visual illusion, we trained a classifier to decode blocks of illusory trials from blocks of trials in which no illusion was perceived. Using leave-one-run-out cross validation, we found that responses in multiple cortical areas were sensitive to the double drift illusion (Fig 2); the classifier could accurately decode illusory drift in all four visual area ROIs (Fig 3A, left).

**Figure 2:**
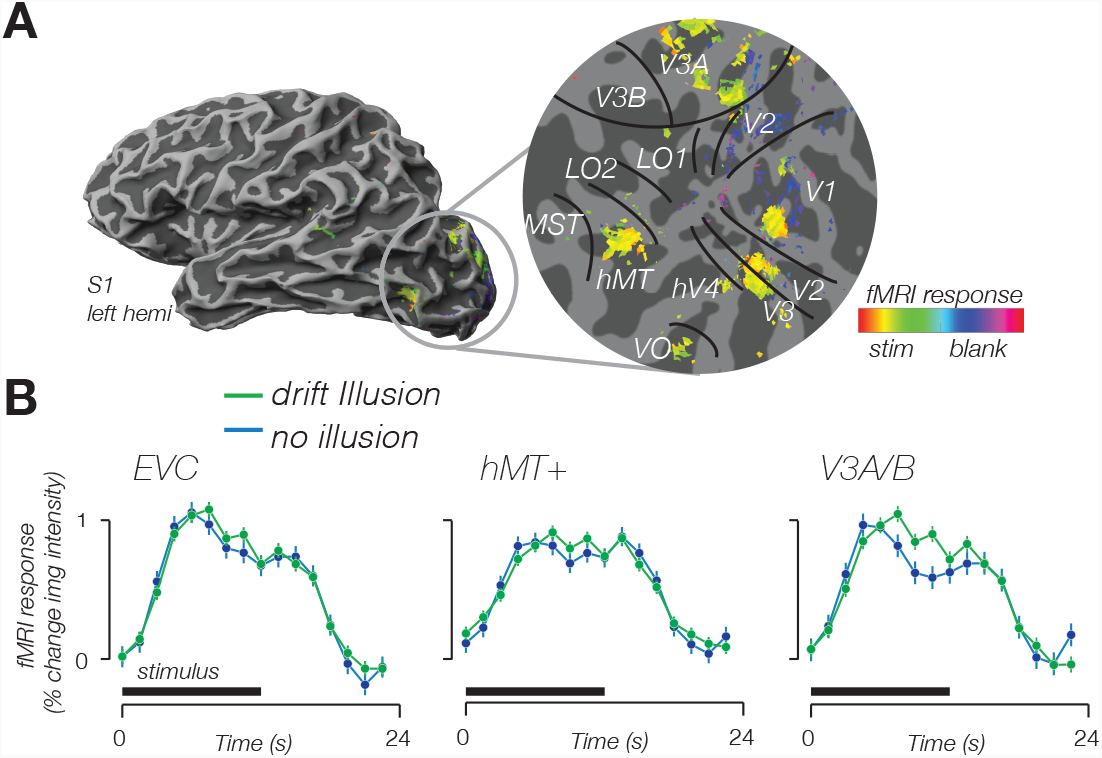
Modulation of fMRI response amplitude during double drift illusion. **Top**, Cortical activity evoked by stimulus localizer. Data from a single participant shown on an inflated cortical surface (left) and a flattened patch of the occipital lobe (right). Boundaries of retinotopic visual areas identified according to an anatomical template. Color indicates the phase of the response. Yellow hues indicate a response in phase with the onset of the stimulus. **Bottom**, Time course of fMRI response from three areas exhibiting a larger response for the double drift illusion than during a control condition that did not produce an illusory drift path.

**Figure 3.**
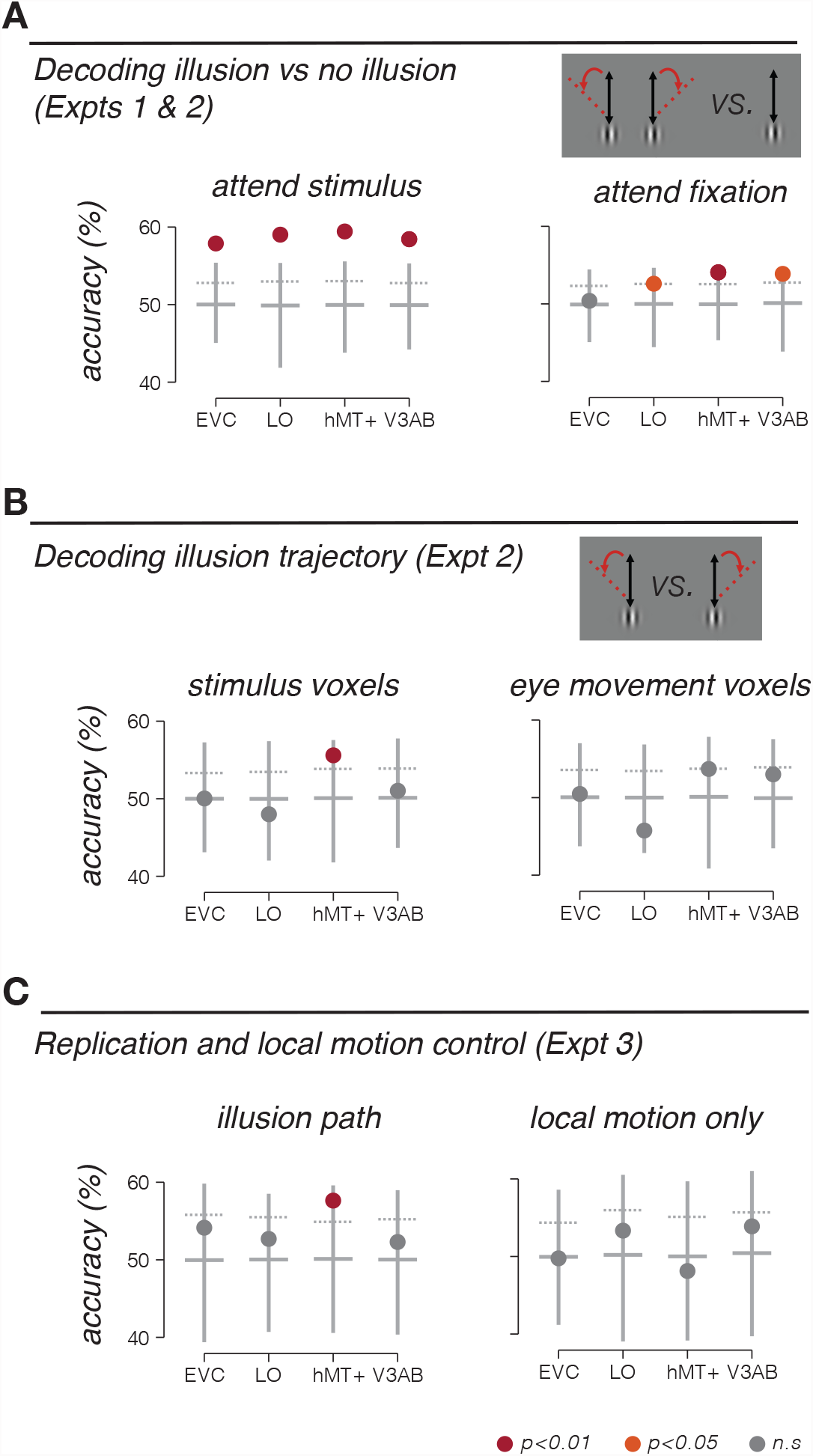
Decoding stimulus drift path. **A**. Accuracy of discriminating the double drift illusion from a control condition that was matched for net motion energy. Participants either attended the peripheral stimulus and reported the presence of the illusion (Expt 1, left), or attended the fovea and reported a luminance decrement at fixation (Expt 2, right). **B**. Accuracy of discriminating rightward vs. leftward drift illusion paths in Expt 2 (attend fixation) based on fMRI responses in voxels selected to match the retinotopic location of the stimulus (left) and voxels selected based on responses to pursuit eye movements (right). **C**. Decoding accuracy for independent replication and control experiments (Expt 3). Left, decoding illusory drift paths, replicating results of Expt 2. Right, decoding local-motion only control conditions, which did not produce a drift illusion. Vertical gray line indicates minimum to maximum range of decoding accuracy obtained with permuted labels. Solid horizontal line, 50^th^ percentile of permuted decoding accuracy; dashed horizontal line, 95^th^ percentile of permuted decoding accuracy. Maroon dot, p<0.01; Orange dot, p<0.05; gray dot, nonsignificant (p>0.05).

**Table 1.** Decoding results for all experiments.

|  |  | EVC | LO | hMT+ | V3A/B |
| --- | --- | --- | --- | --- | --- |
| **Expt 1: Decoding illusion** | Accuracy | 0.5788 | 0.5902 | 0.5943 | 0.5844 |
|  | SD | 0.0734 | 0.0781 | 0.1011 | 0.0626 |
|  | P value | <0.001 | <0.001 | <0.001 | <0.001 |
| **Expt 2: Decoding illusion** | Accuracy | 0.5039 | 0.5260 | 0.5407 | 0.5387 |
|  | SD | 0.0693 | 0.1024 | 0.0788 | 0.0577 |
|  | P value | 0.3800 | 0.0470 | 0.0050 | 0.0110 |
| **Expt 2: Decoding trajectory (stimulus-selective voxels)** | Accuracy | 0.5003 | 0.4800 | 0.5560 | 0.5103 |
|  | SD | 0.0492 | 0.0877 | 0.0726 | 0.0993 |
|  | P value | 0.4930 | 0.8030 | 0.0060 | 0.3380 |
| **Expt 2: Decoding trajectory (pursuit-selective voxels)** | Accuracy | 0.5047 | 0.4575 | 0.5369 | 0.5300 |
|  | SD | 0.0679 | 0.0849 | 0.0815 | 0.0751 |
|  | P value | 0.4160 | 0.9780 | 0.0520 | 0.1000 |
| **Expt 3: Decoding trajectory** | Accuracy | 0.5414 | 0.5269 | 0.5764 | 0.5230 |
|  | SD | 0.0425 | 0.0743 | 0.0526 | 0.0771 |
|  | P value | 0.1140 | 0.1830 | 0.0090 | 0.2480 |
| **Expt 3: Local motion control** | Accuracy | 0.4976 | 0.5333 | 0.4813 | 0.5389 |
|  | SD | 0.0727 | 0.0780 | 0.0905 | 0.0600 |
|  | P value | 0.5250 | 0.1940 | 0.7150 | 0.1330 |

A number of different factors could lead to accurate decoding of the double drift illusion. One possibility is that the decoder was sensitive to neural activity related to computing the location of the stimulus.

Alternatively, it is possible that the perception of the illusion attracted spatial attention, and the classifier was picking up on attentional differences between illusory and non-illusory conditions. To control for this second possibility, we repeated the experiment, but had participants perform a demanding task at fixation that required sustained attention (Haladjian et al., 2018). The fixation task minimized differences in spatial attention to the stimulus across conditions. We again tested whether the classifier could discriminate the double drift illusion from the control condition. While overall decoding accuracy was slightly reduced in this experiment, we found that decoding accuracy remained robust and significant in LO, hMT+, and V3A/B, but not in EVC (Fig 3A, right), consistent with other recent observations (Liu et al., 2019; Ho and Schwarzkopf, 2021). Participants were not attending the stimulus, therefore these results cannot be attributed to differences in spatial attention. Instead, we conclude that the classifier was sensitive to information related to encoding the perceived position of the stimulus during the illusion.

The critical test in this study is whether BOLD fMRI activity in visual cortex can discriminate between different illusory paths. We tested whether a classifier could decode the drift path of the illusion. Of all the visual areas tested, only area hMT+ could discriminate leftward from rightward illusory paths (Fig 3B, left). Because the stimulus remained at a constant retinal location, the ability to discriminate the illusory motion path suggests a non-retinotopic representation of stimulus position.

We next tested two alternative explanations for these results. First, it is conceivable that decoding of the drift path was due to subtle differences in smooth pursuit eye movements, rather than encoding of the stimulus position. While we were not able to record eye movements in the scanner, we did perform a control analysis that argues against this possibility. To test this, we repeated the classification analysis, this time using only voxels that were selective for smooth pursuit eye movements, as identified in a separate pursuit control experiment. In this analysis, we specifically excluded voxels that responded in the stimulus localizer (see *Materials & Methods: Eye movement localizer experiment*). Voxels that responded in the pursuit localizer should be most sensitive to any differences in pursuit eye movements in the main experiment, regardless of whether these voxels are selective for pursuit eye movements themselves, or to the visual consequences of retinal slip during pursuit. But we found that responses in these voxels do not carry information that distinguishes the drift paths, in any of the ROIs (Fig 3B, right). Results from this control analysis suggest that the information being utilized by the classifier is not due to differences in pursuit eye movements.

The stimulus for both illusion drift paths consisted of a combination of a vertical global trajectory and horizontal local motion. Yet rightward and leftward illusion drift paths differed in the order of local motion (see Fig 1). We wondered if the ability of the classifier to discriminate leftward and rightward illusion drift paths could be due to the difference in temporal sequence of events within the trial (e.g., leftward followed by rightward motion). To test this, we scanned another group of participants in an experiment that included both of the illusion conditions from Expt 2, and two control conditions that contained the same local motion, but no global trajectory (and no smooth pursuit eye movements). For the illusory double drift conditions, we again found that drift path was decodable from hMT+, replicating the results from Exp 2 in an independent group of participants (Fig 3C, left). However, the classifier was unable to decode the conditions containing local motion information alone (Fig 3C, right). This result demonstrates that decoding in hMT+ depends on the double drift illusion, and not on the local motion of the stimulus alone.

## Discussion

We found that fMRI BOLD responses in several visual cortical areas could reliably discriminate the double drift illusion from a control condition that was matched for motion energy. In early visual cortex, this result could be explained by attentional effects associated with perceiving the illusion, since when attention was directed away from the illusion decoding in EVC dropped to chance. But in areas hMT+, LO, and V3A/B, decoding persisted even when controlling for spatial attention. Moreover, responses in hMT+ could discriminate the illusory drift path. A number of control experiments indicate that this result cannot be explained by low level stimulus or oculomotor factors. Our results may indicate non-retinal stimulus position encoding in human extrastriate visual cortex.

### Source of illusory drift path information

What is the source of decodable drift path information? One possibility is related to a coarse-scale map for direction of motion, which has been observed throughout visual cortex, including all of the areas included in our study (Wang et al., 2014). The coarse-scale map for direction of motion in early visual areas (V1/V2/V3) is thought to result from an aperture-inward bias: larger responses were observed for motion away from the aperture edge (Wang et al., 2014). In contrast, the coarse-scale map observed in hMT+ did not depend on the aperture boundary, but instead consisted of a bias for motion towards the fovea. Could this fovea-centered bias explain the ability to decode the path of the double drift illusion? In the current study, a fovea-centered bias would predict a leftward preference across voxels within hMT+, since the stimulus was always in the right visual field and leftward motion would be toward the fovea. However, the two illusory conditions (Fig 1A,B) had identical net amounts of leftward and rightward local motion, and identical proportions of time of leftward and rightward illusory drift paths. We think it is hence unlikely that a global motion bias in hMT+ accounts for the observed results.

An alternative account is that differences in BOLD activity to the two illusory drift paths arise because of the topographic organization within hMT+ (Huk et al., 2002; Amano et al., 2009). The rightward drift path begins with illusory drift up-and-to-the-right, which increases the perceived eccentricity of the Gabor (Lisi and Cavanagh, 2015). The path continues with drift down-and-to-the-left, which brings the perceived position back to the original position. This cycle repeats throughout the block of trials. In contrast, the leftward drift path begins with illusory drift up-and-to-the-left, which decreases the perceived eccentricity of the Gabor, and continues with drift down-and-to-the-right, bringing the perceived position back to the original position. Thus, the average eccentricity of the perceived drift path is higher during rightward drift, and lower during leftward drift. This shift in the perceived eccentricity of the stimulus could result in slightly different patterns of activity in hMT+, and this difference could underlie the ability to decode the illusion drift path.

This second account depends on there being an explicit representation of the perceived position of a stimulus in hMT+, while position encoding in EVC is entirely veridical. Since veridical position did not differ for rightward and leftward drift paths, the classifier was unable to decode the drift path from activity in EVC. Consistent with this account, one fMRI study (Maus et al., 2013) has reported that BOLD activity in hMT+ reflects the illusory position during motion-induced position shift. Activity throughout visual cortex is known to encode stimulus position in retinal, not spatiotopic, coordinates (Gardner et al., 2008). However, it is not known if retinotopic coding is also universal in visual cortex for motion illusions. If the second account is accurate our data may imply a difference between EVC and downstream areas in the spatial encoding of illusory motion.

### Spatiotopic coordinates in visual cortex

The double drift illusion results from combining local motion of the Gabor with a global trajectory of the envelope. In the smooth-pursuit variant of the illusion, the envelope only has a trajectory when defined in spatiotopic coordinates, since the stimulus remains at a constant retinal location. The fact that this stimulus condition produces an illusion suggests some degree of spatiotopic processing in the brain. This could be accomplished by the formation of an explicit spatiotopic reference frame (Duhamel et al., 1997; d’Avossa et al., 2007; Crespi et al., 2011). Alternatively this could be accomplished through a computation by which a retinotopic input is combined with an eye position gain field (Merriam et al., 2013). Our data do not speak to which of these two possibilities is more likely.

### Decoding the drift illusion beyond hMT+

In addition to decoding the illusory drift path, patterns of activity in multiple visual areas enabled classification of the perception of the illusion. When subjects were attending the stimulus, illusion decoding was possible in all the visual areas that we studied, raising the possibility that the illusion attracted attention, resulting in a higher BOLD response during illusory blocks (see Fig 2). When subjects were attending to a task at fixation (Exp 2), the illusion could still be decoded from activity in LO, V3A/B, and hMT+, but not from EVC. Previous fMRI studies have claimed that the locus of attention can affect the apparent reference frame in which a stimulus is encoded (Crespi et al., 2011). It is unclear, however, whether attention and task indeed change the spatial reference frame, or instead affect global response amplitudes (Roth et al., 2020), which may constitute an additive signal obfuscating measurement of the underlying reference frame. In Exp 1, subjects performed a task on the stimulus, and so attention was likely directed toward the stimulus. In an earlier fMRI study on the double drift illusion (Liu et al., 2019), subjects also performed a task on the stimulus, and while the tasks were different in the two experiments, in both cases attention was focused on the stimulus. Our results in Exp 2, in which subjects attended fixation, and not the stimulus, demonstrate that attention and task do have an impact on the spatial encoding of the double drift illusion, and highlights the importance of controlling the attentional state of the observer when studying visual reference frames (Crespi et al., 2011).

### Disentangling spatiotopic representations and remapping

Behavioral investigations into mechanisms for visual stability across eye movements have found evidence that both spatiotopic representations and receptive field remapping underly visual stability (Poletti et al., 2013), with the relative contribution of each mechanism depending on the number of intervening saccades. After a single saccade, receptive field remapping is the primary mechanism underlying visual stability, whereas spatiotopic representations become prominent after multiple saccades (Sun and Goldberg, 2016). It is therefore possible that fMRI studies exploring spatiotopic representations were in fact probing retinotopic coding that is updated by remapping across saccades. This is true especially for studies in which stimuli were presented immediately after saccades (McKyton and Zohary, 2007), but also when short periods of time intervene between saccade and stimulus presentation (d’Avossa et al., 2007; Crespi et al., 2011). The version of the double drift illusion employed in the current study did not require saccadic eye movements, making it unlikely that perisaccadic remapping contributed to our results. Rather, our results suggest that stimulus position was encoded in a spatiotopic representation.

## Materials and Methods

### Participants

Data were acquired from 19 healthy participants (11 females, age range 23-34 years, mean 25.8 years) with normal or corrected-to-normal vision. Experiments were conducted with the written consent of each observer. The consent and experimental protocol were in compliance with the safety guidelines for MRI research, and were approved by the Institutional Review Board of the National Institutes of Health. 12 of the 19 participants were scanned in multiple sessions and in multiple experimental conditions. 9, 12, and 5 participants participated in Exp 1, 2, and 3, respectively.

### Stimuli

A Gabor pattern, consisting of a vertically oriented sinusoidal grating (spatial frequency of 1 deg / cycle) within a Gaussian envelope (standard deviation of 1 dva), moved back and forth along a linear, vertical trajectory with a length of 8 deg. The trajectory length was varied slightly (+/– 1 deg) across subjects to accommodate the restricted field of view within the scanner. The Gabor moved according to a linear velocity profile (10 deg/sec) that was smoothed slightly at the top and bottom of the Gabor’s path where it changed direction to facilitate accurate pursuit eye movements. The Gabor’s internal grating moved orthogonally relative to its trajectory with a speed of 6.66 Hz, reversing its direction at the two endpoints of the trajectory. Participants fixated a target (0.2 dva, white dot with a black outer rim) that was positioned 9-12 dva to the left of the Gabor envelope and moved smoothly alongside it. Participants were instructed to pursue the target. Pursuit accuracy was confirmed during the behavioral experiment for each subject prior to the fMRI experiment.

### Pre-scan behavioral experiment

Prior to the first scanning session, participants viewed the double drift stimulus in a behavioral experiment and were asked to judge the angle of the illusion. Participants viewed the illusion in 12 s blocks while pursuing a target that moved smoothly alongside the stimulus. The Gabor completed 8 traversals per block. At the end of each block, the Gabor was replaced with a vertical line, aligned to the physical (vertical) path of the Gabor’s trajectory. Participants manipulated the angle of the line with the keyboard in order to match the perceived path of the Gabor. Participants viewed 25-50 blocks of the illusion.

### Experimental conditions and fMRI design

#### Expt 1: Attend to drift illusion path

The main fMRI experiment consisted of a randomized blocked design with three stimulus conditions, all of which involved the same vertical Gabor envelope trajectory: 1, perceived leftward drift path (internal local motion leftward during upward trajectory; rightward during downward trajectory); 2, perceived rightward drift path (internal local motion rightward during upward trajectory; leftward during downward trajectory); 3, no-illusion control condition (randomized internal local motion, updated at 60 Hz). In all three conditions, participants pursued a fixation dot that moved smoothly and predictably alongside the Gabor, so that the Gabor remained at a constant retinal location throughout the experiment. Under conditions 1 & 2, the Gabor’s perceived drift path differed from its actual trajectory by several degrees. In condition 3, the Gabor’s perceived path matched its actual trajectory (i.e., there was no illusion). Each of the three conditions contained the same global and local motion energy and required the same smooth pursuit eye movements.

Each Gabor traversal lasted for 1.5 s (750 ms up, 750 ms down). The Gabor completed 8 traversals in a 12 s block. The three conditions were randomly interleaved, and experimental blocks alternated with 12 s blocks of fixation in which both the pursuit target and the Gabor remained stationary, with no internal motion. Participants were instructed to press one of three buttons at the end of each experimental block indicating the direction of the illusion (‘1’ for leftward drift illusion, ‘2’ for no illusion, and ‘3’ for rightward drift). The double-drift illusion is typically strong and unambiguous, even during smooth pursuit eye movements. Accordingly, participants performed the attend-to-stimulus task with nearly 100% accuracy. Each fMRI run lasted for 288 s and included 4 blocks of each of the 3 conditions in a randomized order.

#### Expt 2: Attend to fixation target

Experimental design and stimuli were identical to Expt 1, except for additional luminance decrements of the fixation target. Participants’ task was to press a button when they detected the brief (250 ms) luminance decrement. Luminance decrements were determined using an adaptive, 1-up-2-down staircase procedure (Levitt, 1971), producing a detection rate of approximately 70%. The Gabor stimulus was not relevant to the task and subjects were not instructed to attend to it. Each fMRI run lasted for 288 s and included 4 blocks of each of the 3 conditions in a randomized order.

#### Expt 3: Local motion control

This experiment controlled for differences in the pattern of local internal motion in Expts 1 and 2. While the two illusory drift paths (leftward drift, rightward drift) were balanced for net local and global motion energy, they differed in the order of motion direction. In the leftward drift illusion condition, local motion started leftward (for 750 ms) and was followed by rightward motion (for 750 ms). Vice versa for the rightward drift illusion condition. Expt 3 consisted of 4 conditions.

Conditions 1 and 2 were identical to the two illusory conditions in Expts 1 and 2. However, in this experiment there were two additional control conditions in which participants viewed the same patterns of local motion as in conditions 1 and 2, but the Gabor did not move across the screen (no global motion trajectory), nor were there smooth pursuit eye movements. Instead, participants fixated a stationary target alongside a stationary Gabor containing internal motion to the left and right. In condition 3, the order of local motion was the same as in condition 1 (leftward followed by rightward). Condition 4 matched the local motion of condition 2. Participants were instructed to press a button when they detected the brief (250 ms) luminance decrement. Each fMRI run lasted for 288 s and included 3 blocks of each of the 4 conditions in randomized order.

#### Stimulus-only localizer experiment

In each scanning session, participants were scanned in a stimulus-only localizer experiment in which the stimulus appeared and disappeared in a two-condition block alternation protocol (9 s on, 9 s off; 14 blocks per fMRI run, lasting 252 s). Participants maintained fixation on a stationary target. A vertical Gabor stimulus appeared at the same size and eccentricity as in the main experiment. The Gabor contained internal local motion that changed direction randomly every 250 ms. After 9 s, the stimulus disappeared and participants continued to fixate. Three stimulus-only runs were included in each scanning session, one run at the beginning of the session, one in the middle, and a third run at the end of the session. Participants did not perform a behavioral task during the stimulus-only experiment.

#### Eye movement localizer experiment

In each scanning session, participants were scanned in an eye movement localizer experiment in which participants tracked a moving fixation dot that was identical to the double drift illusion experiments, except that there was no peripheral Gabor stimulus. Responses to pursuit eye movements were measured in a two-condition block alternation protocol (9 s pursuit, 9 s fixation; 14 blocks per fMRI run, each lasting 252 s). Two eye movement localizer runs were included in each scanning session, one run at the beginning of the session and a second run at the end of the session. Participants did not perform a behavioral task during eye movement localizer experiment.

### Experimental setup - behavioral

Stimuli were generated using Matlab (MathWorks, MA) and MGL (Gardner et al., 2018) on a Macintosh computer, and presented on a 61-inch screen (BenQ XL242OZ) positioned 57 cm away from the participant. Participants were seated in a darkened room and were head stabilized by a chin rest. An Eyelink 1000 eye-tracking system was used to measure binocular eye position at 1000 Hz. Eye-tracking calibration was performed at the beginning of the session and repeated intermittently throughout the session to ensure that eye tracking accuracy remained within 1 deg of visual angle throughout the experiment.

### Experimental setup -fMRI

Stimuli were generated using Matlab (MathWorks, MA) and MGL (Gardner et al., 2018) on a Macintosh computer. Stimuli were displayed via a PLUS U2-1200 LCD projector (resolution: 1024 × 768 pixels; refresh rate: 60 Hz) onto a back-projection screen in the bore of the magnet. Participants viewed the display through an angled mirror at a viewing distance of approximately 58 cm, producing a field of view of 20.5 deg × 16.1 deg. fMRI data were acquired from participants on a research-dedicated Siemens 7T Magnetom scanner using a 32-channel head coil, located in the Clinical Research Center on the National Institutes of Health campus (Bethesda, MD). Functional imaging was conducted with 56 slices oriented parallel to the calcarine sulcus covering the posterior half of the brain: TR: 1500 ms; TE 23 ms; FA: 55°; voxel size: 1.2 × 1.2 × 1.2 mm with 10% gap between slices; grid size: 160 × 160 voxels. Multiband factor 2, GRAPPA/iPAT factor 3. The slices covered all of the occipital and parietal lobes, and the posterior portion of the temporal lobe.

For each participant, a high-resolution anatomy of the entire brain was acquired by co-registering and averaging between 2 and 8 T1-weighted anatomical volumes (magnetization-prepared rapid-acquisition gradient echo, or MP2RAGE; TR: 2500 ms; TE: 3.93 ms; FA: 8°; voxel size: 0.7 × 0.7 × 0.7 mm; grid size: 256 × 256 voxels). The averaged anatomical volume was used for co-registration across scanning sessions and for gray-matter segmentation and cortical flattening. Functional scans were acquired using T2*-weighted, gradient recalled echo-planar imaging to measure blood oxygen level-dependent (BOLD) changes in image intensity (Ogawa et al., 1990). The inplane anatomical was aligned to the high-resolution anatomical volume using a robust image registration algorithm (Nestares and Heeger, 2000).

Prior to the first experimental functional run of each session, 30 volumes were acquired with identical scanning parameters and slice prescription as the subsequent functional runs, except for the phase encoding direction which was reversed. This single reverse phase-encoded run was used to estimate the susceptibility-induced off-resonance field using a method similar to that described in (Andersson et al., 2003) as implemented in FSL (Smith et al., 2004). This estimate was then used to correct the spatial distortions in each subsequent run in the session.

### fMRI preprocessing and analysis

The anatomical volume acquired in each scanning session was aligned to the high-resolution anatomical volume of the same participant’s brain, using a robust image registration algorithm (Nestares and Heeger, 2000). Head movement within and across scans was compensated using standard procedures (Nestares and Heeger, 2000). The time-series from each voxel was divided by its mean to convert from arbitrary intensity units to percent modulation and high-pass filtered (cutoff = 0.01 Hz) to remove low-frequency noise and drift (Smith et al., 1999).

### ROI definition

Regions of interest were defined according to an anatomical templated (Benson and Winawer, 2018). Because of the inherent imprecision of such a template, we combined nearby regions to form larger ROIs which we analyzed. V1, V2, and V3 were combined to form an early visual cortex ROI (EVC); LO1 and LO2 were combined into LO; V3A and V3B were combined into V3A/B; TO1 and TO2 were combined to form hMT+, corresponding to MT and MST.

### Data analysis for stimulus-only and eye-movement localizers

The three stimulus localizer scans were averaged together, and the two eye-movement scans were averaged together. The first cycle of each averaged time series was discarded, leaving 13 cycles. Each individual voxel’s time course was fitted to a cosine with a period matching the cycle duration of 12 volumes (18 s). Each voxel was then assigned the correlation coefficient and phase of the best-fitting cosine

### GLM analysis

BOLD fMRI timeseries were averaged across all voxels within each ROI. Trials were then divided into 2 conditions: double drift illusion (either leftward or rightward) and no illusion. Each condition was modeled by 16 predictors, one for each time point in the 24 s following the beginning of the trial.

Deconvolution was performed by multiplying the pseudoinverse of the condition predictor matrix with the time series (Dale, 1999). This procedure yields two hemodynamic response functions for each ROI. The average of the 2 response functions was used as the ROI’s hemodynamic response function for the next analysis step. Next, a design matrix was constructed for each ROI, with a single HRF function modeling each 12 s block. Response amplitudes were then computed by taking the pseudoinverse of this design matrix and multiplying it with the single voxel timeseries.

The goal of this analysis was to estimate a response amplitude for each voxel on each scanning run. The first step was to estimate a single hemodynamic response function for each ROI and each participant. This was accomplished by averaging across all voxels within the ROI, and then using deconvolution (Gardner et al., 2008) to estimate a hemodynamic response function (collapsing across the different conditions) over a 24 s period following the beginning of the block. Next, this single hemodynamic response was used to create a design matrix, treating each of the conditions independently. Response amplitudes were then computed by taking the pseudoinverse of this design matrix and multiplying it with the timeseries for each individual voxel within the ROI. This procedure allowed for differences in the shape of the hemodynamic response across different ROIs and across different participants.

### Decoding analysis

In multivariate classification analysis of fMRI data, each condition is represented by a set of points in multi-dimensional space, with dimensionality equal to the number of voxels and each point corresponding to a single measurement. Accurate decoding is possible when the responses corresponding to different conditions form distinct clusters within this high-dimensional space. We measured the amplitude of the fMRI response during 12 s blocks of trials, in which each block consisted of 8 up-down traversals of the double drift illusion. We took the beta weight from the GLM analysis (see above) as the amplitude of the response during each 12 s block as a single input to the classification analysis. These response amplitudes were stacked across blocks within a run, and across runs within a session, forming an m × n matrix, with m being the number of voxels in the region of interest and n being the number of repeated measurements in the session. The value of n was typically 64 (for 14 of 19 participants), and ranged from 48 (1 participant) to 96 (4 participants). Decoding was performed with a maximum likelihood classifier, using the Matlab function “classify” with the option “diagLinear” (Roth et al., 2018). Decoding accuracy was computed using leave-one-run out cross-validation. The m × n data matrix was partitioned along the n dimension (repeated measurements) into training and testing sets, in which the training set consisted of the blocks from all but one of the runs, and the testing set included the blocks (from all three conditions) from the left out run. Because the data in the training and testing sets were drawn from different runs in the same session, they were statistically independent. The training set was used to estimate the parameters (multivariate means and variances) of the maximum-likelihood classifier. The testing set was then used for decoding. Decoding accuracy was determined as the proportion of the test examples that the classifier was able to correctly assign to one of the two illusory drift paths.

The leave-one-run-out cross validation procedure resulted in a single decoding accuracy estimate per ROI per session. A non-parametric permutation test was used to evaluate the significance of this decoding accuracy. Specifically, we constructed a distribution of accuracies expected under the null hypothesis that there is no difference between the two illusory drift paths. To generate a null distribution decoding accuracy, we permuted the block labels for each run, and repeated the leave-one-run-out decoding analysis. Repeating this randomization 1,000 times yielded a distribution of accuracies expected under the null hypothesis. Accuracies computed using the unrandomized training data were then considered statistically significant when decoding accuracy was higher than the 95^th^ percentile of the null distribution (P < 0.05, one-tailed permutation test).

## Acknowledgements

Supported by the Intramural Research Program of the National Institutes of Health (ZIAMH002966).

